# The Threshold Elemental Ratio of an ectotherm decreases then increases with rising temperature

**DOI:** 10.1101/681239

**Authors:** Thomas Ruiz, Apostolos-Manuel Koussoroplis, Michael Danger, Jean-Pierre Aguer, Nicole Morel-Desrosiers, Alexandre Bec

## Abstract

Earth is currently facing unprecedented global changes, hurrying scientists to provide predictive tools to explore the futures responses of ecosystems. Among those changes, temperature increase and alterations of nutrient availabilities largely drive consumer performances, yet their interactive effect remains poorly understood. Here we investigate how the dietary C:P ratio that optimizes consumer growth (TER_C:P_: Threshold Elemental Ratio) changes along temperature gradients by combining a TER_C:P_ model and growth experiments on the model organism *Daphnia magna*. Both lines of evidence show that the TER_C:P_ responds to temperature in an U-shaped fashion. This shape reconciles previous contradictive observations into a common framework, thereby improving our capacity to forecast the combined effects of nutrient cycle and climatic alterations on ectotherms.

## Main Text

Nutrition and temperature are two central drivers of ectothermic animal performances. Under the influence of global change, both food quality (*1*) and temperature (*2*) are dramatically affected, inducing important constraints on ectotherm populations (*3*). While the effects of each factor independently are well studied, there is a stringent need to determine how nutrition and temperature interactively affect consumer biomass production and the resulting consequences on ecosystem functioning (*4, 5*).

This study shows how the dietary stoichiometry that optimizes consumer growth changes along temperature gradients. A well-established framework for understanding how biomass production responds to variation in food stoichiometry is the threshold elemental ratio (TER) (*6, 7*). The TER is the ratio at which nutrients need to be present in food in order to maximize consumer performance (*6*). A special emphasis has been given to the TER relating carbon (C) to phosphorus (TER_C:P_) or nitrogen (TER_C:N_) since they separate an energy-limitation (C-limitation) of growth from a nutrient-limitation of growth (*7*). Deviation of food stoichiometry from this optimal ratios is viewed as a decrease of quality since it reduces consumer growth rate (*7*). Such an imbalance between TER and food stoichiometry can also alter P and N recycling in ecosystems because consumer is forced to actively release the nutrient in excess relative to its needs (*8*). Many studies have stressed the importance of interspecific differences in the TER to explain ecological processes (*6, 9*). Yet, the TER is suspected to change intra-specifically depending on environmental conditions (*5*). Kindled by the ongoing global temperature increase and alterations of nutrients cycles (*2*), a debate is emerging on whether and how the TER of species could be affected by temperature (*10*–*12*). Based on indirect experimental evidence and different lines of argumentation, some predict TER increases with increasing temperature (*10, 13*) while others predict the opposite (*12, 14*). This absence of consensus clearly hinders our ability to predict the consequences of climate changes on matter and energy flows within ecosystems (*5*).

Here, we address this issue by combining a TER model and a unique dataset of *Daphnia magna* TER_C:P_ measured along a wide range of temperatures. Our model differs from previous approaches (*15*) in that all processes involved in the determination of the TER are hypothesized to be temperature-sensitive **(Fig.1)**. We assume that certain physiological processes such as ingestion rate and assimilation efficiency respond to temperature with typical Thermal Performance Curves (TPC) (*16, 17*) while others such as metabolic rate display an exponential response. Contrary to previous models predicting monotonic changes of the TER with temperature (*15*), our model suggests that the TER_C:P_ responds non-monotonically to temperature (**Fig.1**). In our model, the resulting reaction norm of the TER to temperature is “U-shaped”: The TER initially decreases with increasing temperature until a minimal value, hereafter named *Threshold Minimising Temperature* (TMT). At the TMT, the dietary C requirements relative to those of P are minimized. Above the TMT, the TER increases again with temperature indicating a rise in the relative C demand. This prediction is supported by the somatic growth rate measurements on *Daphnia* exposed to a factorial combination of dietary C:P ratios and temperatures (**Fig.2 A**). At 18°C the boostrapped TER_C:P_ of 339 (95% CI [315, 360]) decreases down to a minimum value of 194 (95% CI [161, 218]) at 22°C.This temperature can be considered as our *Daphnia* clone’s TMT since TER increases again above it up to 400 (95% CI [395, 405]) at 28°C (**Fig.2 B).**

**Figure 1:**
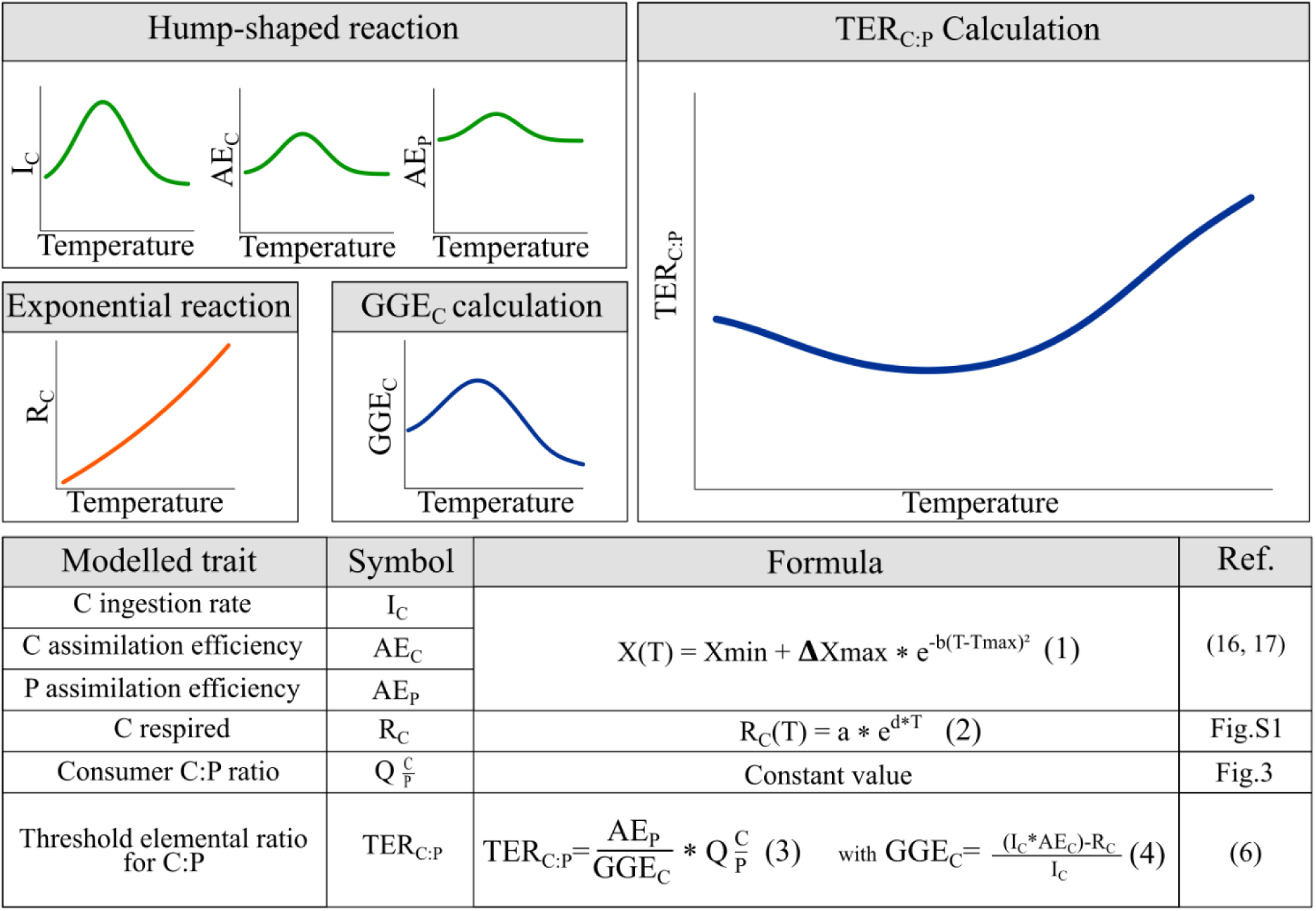
Model of the TER_C:P_ temperature reaction norm. Thermal performance curves are assigned to each process underlying the TER_C:P_. C Ingestion rate (Ic) and assimilation efficiency (AEc) as well as phosphorus assimilation efficiency (AEp) are modelled using equation (1), where X is the trait of interest, **Δ**Xmax the difference between minimum (Xmin) and maximum trait value. T is temperature, Tmax the optimal temperature and *b* is a coefficient determining the decrease rate around Tmax. Respiration (Rc) is modelled using equation (2) with *a* and *d* coefficients being the proportionality constant and the scaling exponent, respectively. The consumer C:P ratio (Q_C:P_) remains unchanged with temperature.

**Figure 2:**
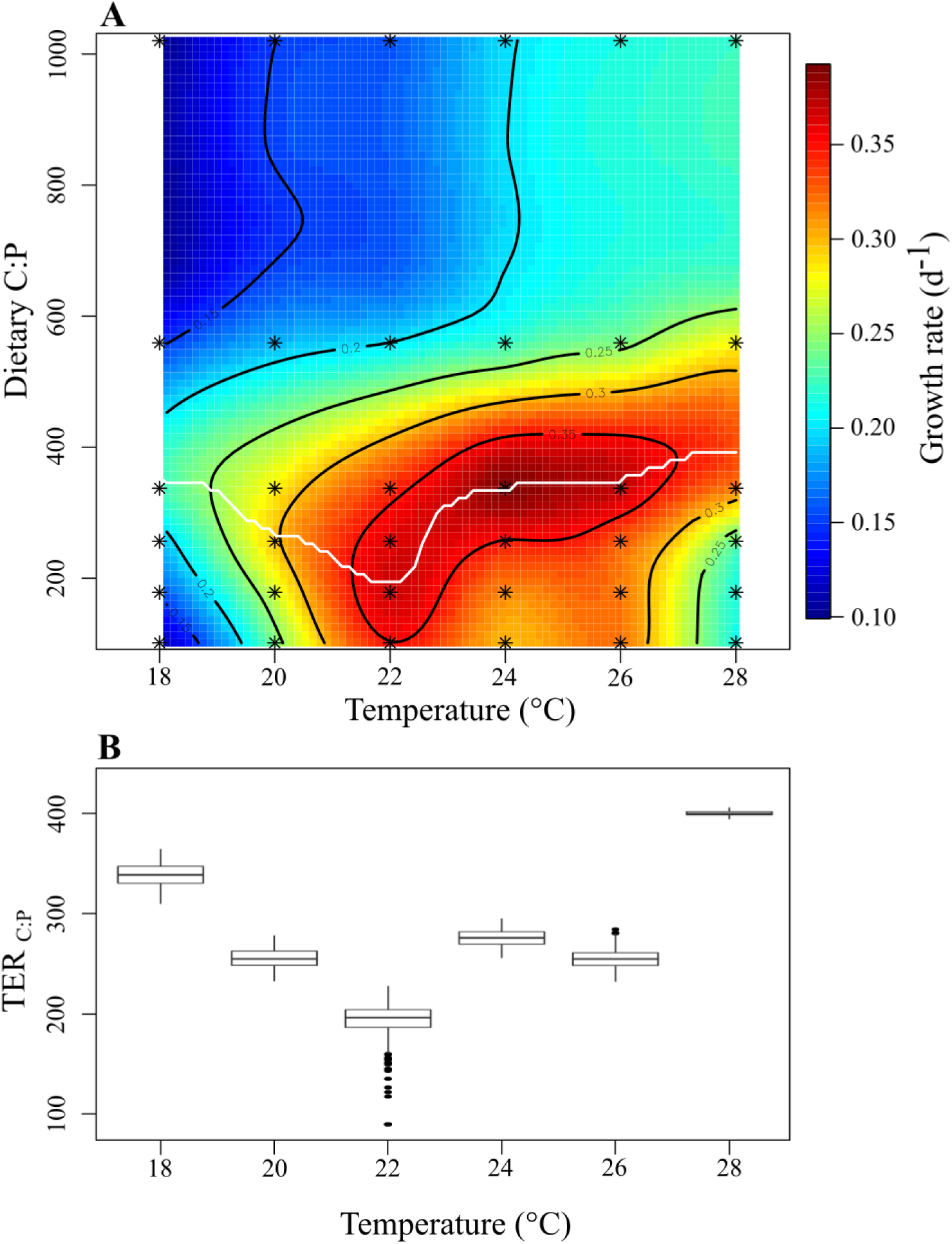
**A/** Growth rate of *Daphnia magna* exposed to a factorial combination of dietary molar C:P ratio and temperature. The white line represents the maximum growth rate reached at each temperature (i.e. TER_C:P_). Asterisks are the experimentally determined values. 3D version available **(Fig.S2).** **B/** Bootstraped TER_C:P_ of daphnids versus temperature. Non-overlapping TER values between temperatures show significant differences. The black points are outliers.

Our findings suggest that previous apparent contradictory predictions (*10, 12*–*14*) can be reconciled within a single framework. For a given species, both temperature-driven decreases and increases of the TER are possible, depending on the range within which temperature changes. According to Cross *et al.* (2015) (*5*) the prediction of TER increase with temperature is based on two arguments. The first argument is that the somatic C:P in consumer tissues should increase at higher temperatures which, holding all other terms in equation 3 **(Fig.1)** constant, should lead to an increase in TER. This argument is based on a general latitudinal trend of conspecific populations with long-term adaptation to their local temperatures (*18*). However, this might not hold for the temperature variability experienced by a single population over shorter temporal scales. Our data shows that there is no significant changes in the somatic C:P ratio neither across dietary (F_(5,124)_=0.763, p=0.384) or temperature (F_(5,124)_=0.139, p=0.710) treatments nor due to their interaction (F_(5,124)_=1.553, p=0.215) **(Fig.3).** The second argument is that the C gross growth efficiency (GGEc) should generally decline with temperature which should also lead to a rise of the TER with temperature. However, this assumption of a monotonic decline of GGEc with temperature is likely unjustified. As experimentally demonstrated in various instances, the ingestion rate of an ectotherm follows a typical TPC (*16, 19*). Our model shows that the asymmetry in temperature-scaling between C ingestion and C respiration yields a hump-shaped thermal reaction norm of the GGEc.

**Figure 3:**
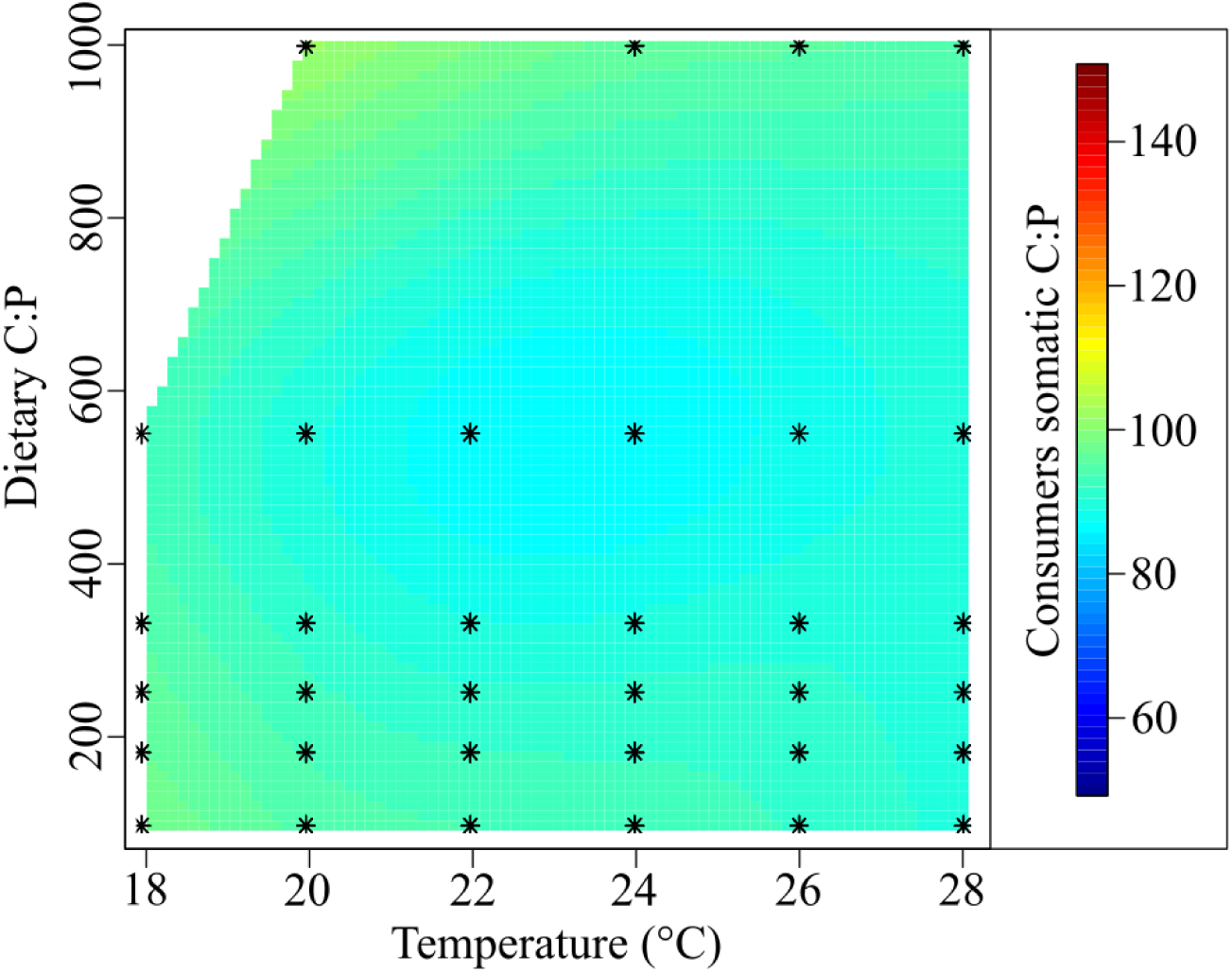
Somatic C:P ratio of *Daphnia magna* exposed to a factorial combination of dietary molar C:P ratio and temperature. Asterisks are the measured points. The somatic C:P was not affected by the dietary C:P (F_(5,124)_=0.763, p=0.384), temperature (F_(5,124)_=0.139, p=0.710) or their interaction (F_(5,124)_=1.553, p=0.215).

Our results indicate that thermal and stoichiometric traits of ectotherms could be linked through the thermal reaction norm of the GGEc. As other ectotherm physiological rates (*20, 21*) the thermal reaction norm of GGEc could also be submitted to physiological trade-offs. For example, reaching a high GGEc peak performance could come at the cost of a narrow thermal performance range. Conversely, maintaining a less variable GGEc over a broader temperature range could imply a reduced peak performance, a situation broadly known as the specialist-generalist trade-off (*20*). Our model predicts that the high peak GGEc of a specialist at optimal temperature also yields a substantial reduction of the TER **(Fig. 4)**. In other words, the thermal specialist potential for higher performances may require a larger amount of P to be reached as compared to thermal generalist. The same reasoning could apply to other commonly observed patterns in ectotherm thermal traits. For example, thermodynamic constraints on enzyme kinetics could imply that for the same thermal breadth, warm-adapted organisms might be able to achieve higher peak GGEc than cold adapted ones (“hotter is better” hypothesis, (*22*)). As in the case of thermal specialists discussed above, warm adapted organisms should have higher P requirements at their optimal temperature than cold adapted ones. Interestingly, this prediction resembles that of the growth rate hypothesis (GRH - (*23*)) which states that sustaining higher growth rates requires a higher P supply. The GRH is based on the observation that taxa with high growth rate potential have higher P body content. Here we show that with a constant body P as in our case, this positive relationship between growth rate potential and P requirements arises from the thermal performance curve of the GGEc. Overall our study highlights the necessity to explore in more depth the links between thermal and stoichiometric traits to predict individual performances in a warming environment. The thermal response of TER we show here constitutes an important step towards unifying thermal ecology and ecological stoichiometry which would enhance our ability to anticipate populations and community dynamics in response to global change. (*5, 24, 25*).

**Figure 4:**
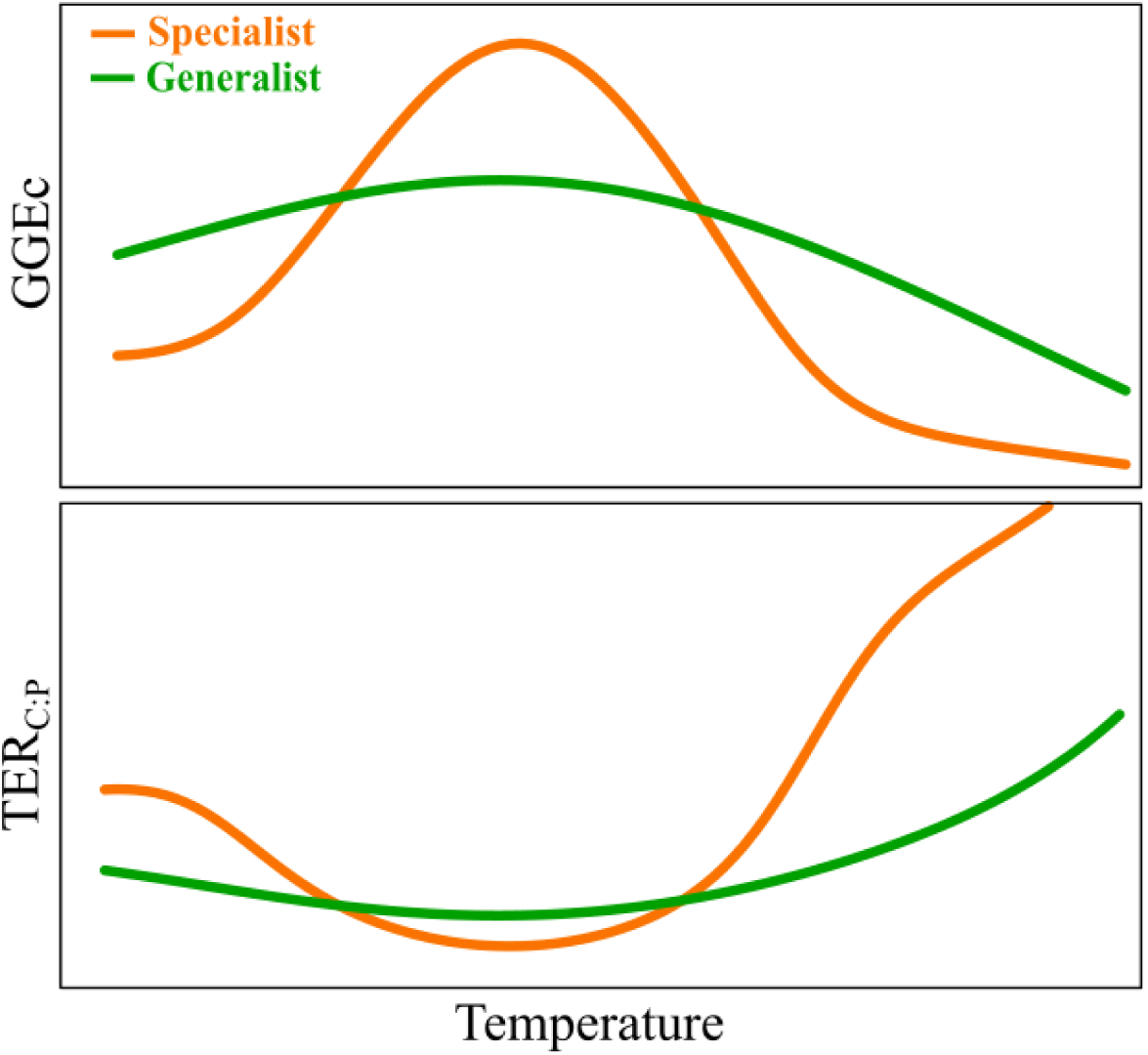
Hypothetical specialist (orange) and generalist (green) thermal performance curves for consumer C gross growth efficiency (GGEc) and the resulting TER_C:P_.

## Acknowledgments

We thank Marie-Elodie Perga for revising an earlier draft version of this manuscript.

## Author contributions

AB, AMK, MD and TR conceived the idea and developed the experimental protocol. TR ran the experiment and acquired the data. AMK and TR developed the model and statistical analysis. TR, AB and AMK led the writing of the manuscript. All authors contributed critically to the draft and gave the final approval for publication.

## Data and materials availability

In case of acceptance of the manuscript, data supporting the paper conclusions and R scripts associated will be deposited on Figshare and the link will be included here.

## Materials and Methods

### Theory

Our model relies on the formulation of the TER by Frost et al. 2006 (*6*) but conversely to previous works we define a reaction norm for each underlying process involved in the determination of the TER. Since virtually all biological processes are enzyme-based, their reaction norms to temperature present a temperature optimum below and above which process rate decreases (*26*). However, in some cases the reaction norm can be simplified into a monotonic function. Accordingly, the thermal reaction norm of respiration is simplified using an exponential function (**Equation 2**) (*27*). This simplification is valid since experimental evidence suggests that metabolic rate, a proxy of respiratory rate, increases with temperature up to temperatures that the organism never experiences in nature (**Fig.S1**). The metabolic rate may also change following dietary stoichiometry (*28*), however we have chosen to ignore this process in order to keep our model as simple as possible. We assume that all other physiological rates and efficiencies that determine the TER exhibit a non-monotonic response to temperature within our thermal ranges. Because it involves biologically tractable parameters, we used a modified Gaussian:

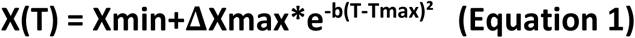

where X is the trait of interest, **Δ**Xmax the difference between minimum (Xmin) and maximum trait value. T is temperature, Tmax the optimal temperature and *b* is a coefficient determining the decrease rate around Tmax. Carbon and nutrient assimilation efficiency (AE) are typically considered to be constant in TER models (*15*). However to the best of our knowledge, there is no foundation to such an assumption. Rather, experimental data on various ectotherms suggest a variation of assimilation efficiency with temperature including both increase (*29*) and decrease (*30*) at higher temperatures. Here, we argue that these diverse responses could be partial observations of an unique underlying hump-shaped response (*17*). We therefore model the thermal performance curves of *AE*_*c*_ and *AE*_*p*_ with the same way as for carbon ingestion rate **(Equation 1)**. This assumption increases the curvature of the TER thermal reaction norm but does not fundamentally change predictions (See sensitivity analyses below).

To our knowledge, the combined effect of temperature and dietary C:P on consumer somatic C:P remains unknown. Hence, we measured the somatic C:P of *Daphnia* exposed to a factorial combination of dietary C:P ratios and temperatures. Our results show that, for our *Daphnia* clone, neither dietary C:P nor temperature significantly affects consumers somatic C:P. We therefore kept C:P constant in the model.

### Sensitivity analysis

To evaluate the robustness of our model predictions, we conducted a sensitivity analysis. The analysis reveal that thermal performance curve for GGE_C_ is the main determinant of the TER_C:P_ shape **(Fig.S3)**. As long as AE_P_ remains higher than AE_C_ (*31*), assuming a flat AE_P_ thermal reaction norm has only a marginal effect on the shape of the TER_C:P_ thermal reaction norm **(Fig. S4)**. The C ingestion rate and assimilation efficiency are the main drivers of the thermal reaction norm of GGE_C_. When AEc remains constant, the shape of GGE_C_ thermal reaction respond to the TPC of C ingestion **(Fig. S5)**. C respiration as a marginal effect on the shape of the GGE_C_ thermal reaction confined within the higher temperature ranges **(Fig. S7)**. As a result, the global shape of TER is primarily condition by C acquisition (i.e. Ic and AEc).

### *Daphnia* maintenance

We tested our model predictions using a clonal line of *Daphnia magna* as a model organism. Daphnids were kept in Volvic water© at 22°C on a 14:10h day:night cycle and fed with *Chlamydomonas reinhardtii* (C:P ratio 331, 3mgC.L^−1^) for >3 generations. Daphnia culture medium was renewed daily to keep optimal feeding conditions and water quality.

### Preparation of food suspensions

*Chlamydomonas reinhardtii* was used as food in all experiment. Algae were grown on modified WC medium with 14.2mg.L-1, 7.1 mg.L^−1^ or 0.35 mg.L^−1^ phosphates (PO_4_^3−^) providing three cultures with respectively a high, intermediate or low content in phosphorus. When enough algal biomass was available for the experiment, cultures were stopped, concentrated and freeze-dried. Freeze-drying process minimize algae denaturation (*32*) and ensure that dietary treatments remain similar during all the experiment. Molar C/P ratios of freeze-dried algae were 95 (high P), 331 (intermediate P) and 1014 (low P). During the growth experiment, food suspensions were daily prepared by mixing freeze-dried algae stocks diluted in Volvic© water (**Table S1**).

### Experimental procedure

To provide a factorial combination of dietary C:P and temperature we performed growth experiments on *Daphnia magna* fed on 6 stoichiometric treatments (with C:P ratios of 95 – 172 – 250 – 331 – 553 – 1014) at 6 different temperatures (18-20-22-24-26 and 28°C). For each temperature, the growth experiment was performed with a strictly similar protocol. For the experiments, 8h old neonates obtained from the 3^rd^ or 4^th^ clutch were randomly distributed in 2 replicate glass jars (250mL, 12 ind./jar) per dietary treatments. The culture medium composed of Volvic© water with *ad libitum* food (6mg dry weight.L^−1^) was renewed daily to maintain optimal feeding conditions during all the experiment. Individuals were raised until they reached maturity, then dried 48h at 60°C, weighed (Sartorius ME36S; Goettingen-Germany: precision 0.001mg) and stored before elemental analysis (*see* below). The average growth rate in each jars was calculated as follow:

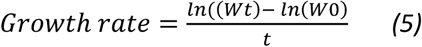

where W_t_ is the average weight (in µg) at age of maturity t (in days) and W_0_ the average weight (in µg) of neonates. Dried daphnids were then used to determine their C:P molar ratio. Carbon was determined using a CHN analyser (NA 2100 Protein, ThermoQuest CE Instruments, Milano, Italy)) and phosphorus content was estimated using a colorimetric determination based on potassium persulfate method (*28*). At maturity, a subset of daphnids from the C:P 250 treatment were used to estimate their metabolic rate. The metabolic rates obtained by microcalorimetry (*28*), were mass-corrected using a 0.75 metabolic-mass scaling exponent (*33*) and plotted against temperature to determine the temperature dependence of metabolism (**Fig. S1).**

### Statistical analyses

For each temperature the confidence intervals around the TER values were generated by nonparametric bootstrapping (*34*). Briefly, for each temperature the growth versus C:P data was resampled a thousand times. For each iteration, we applied the following modified Gaussian function (*26*)(equation 6)

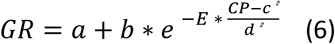

with ***a*** the minimum growth rate on the observation range, ***E*** a constant normalization parameter, ***CP*** the C:P ratio of resources. ***b*** represent the difference between minimum growth rate (***a***) and maximum growth rate. ***d*** constraints the shape of the curve around the optimal value and ***c*** is the estimated TER. **b, c** and **d** are automatically adjusted by non-linear regression to optimize the fit between data and function. The dietary C:P that maximized growth rate was considered to be the TER. The thousand estimated TER values were used to generate the 95% confidence interval of the TER at each temperature (**Fig. 2 B**). The C/P molar ratios of daphnids were compared using Two-way ANOVA after a log-transformation of the data to satisfy ANOVA assumptions. All statistical analysis and model estimation were performed using the software R v.3.4.3 (R Core Team 2018) with an alpha error set at 0.05.

---

**Figure S1:**
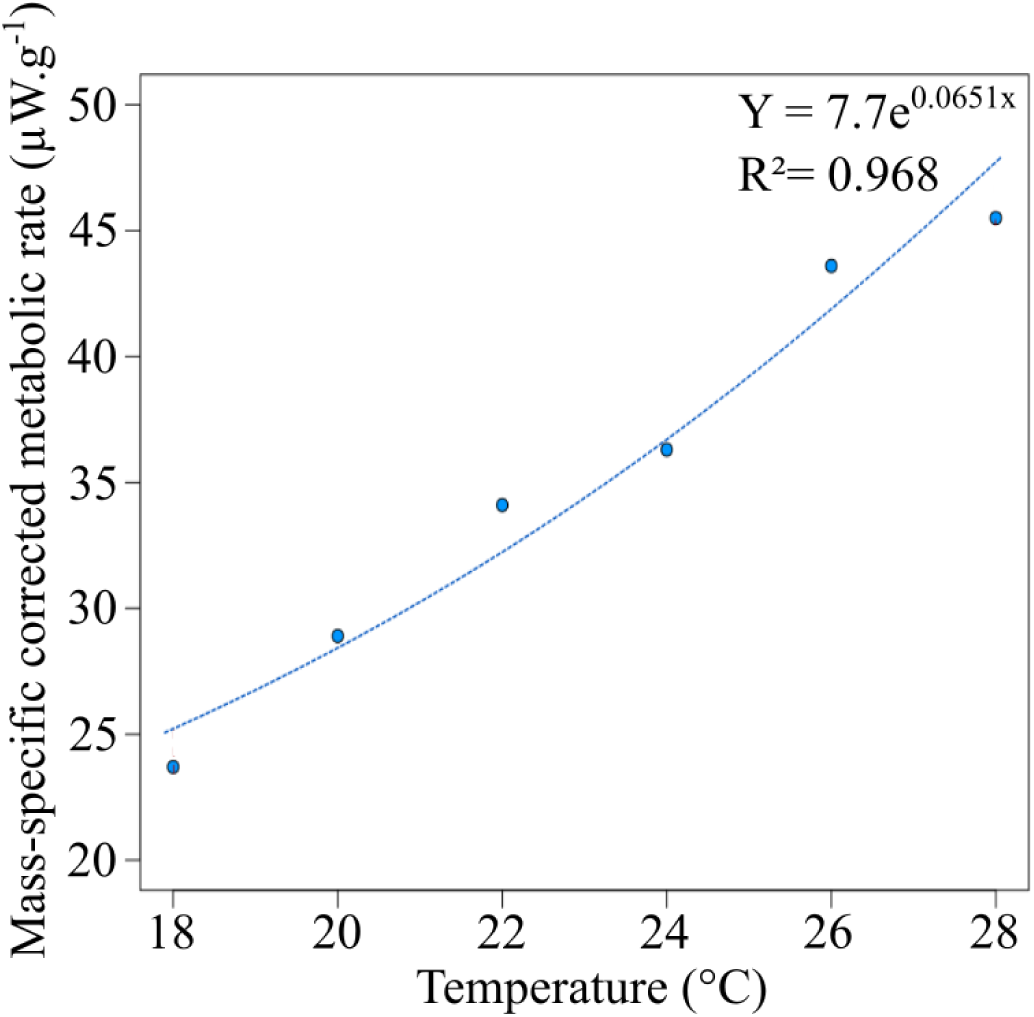
Mass specific metabolic rate versus temperature. Metabolic rate is corrected to 50µg using a 0.75 metabolic-mass scaling exponent (*33*).

**Figure S2:**
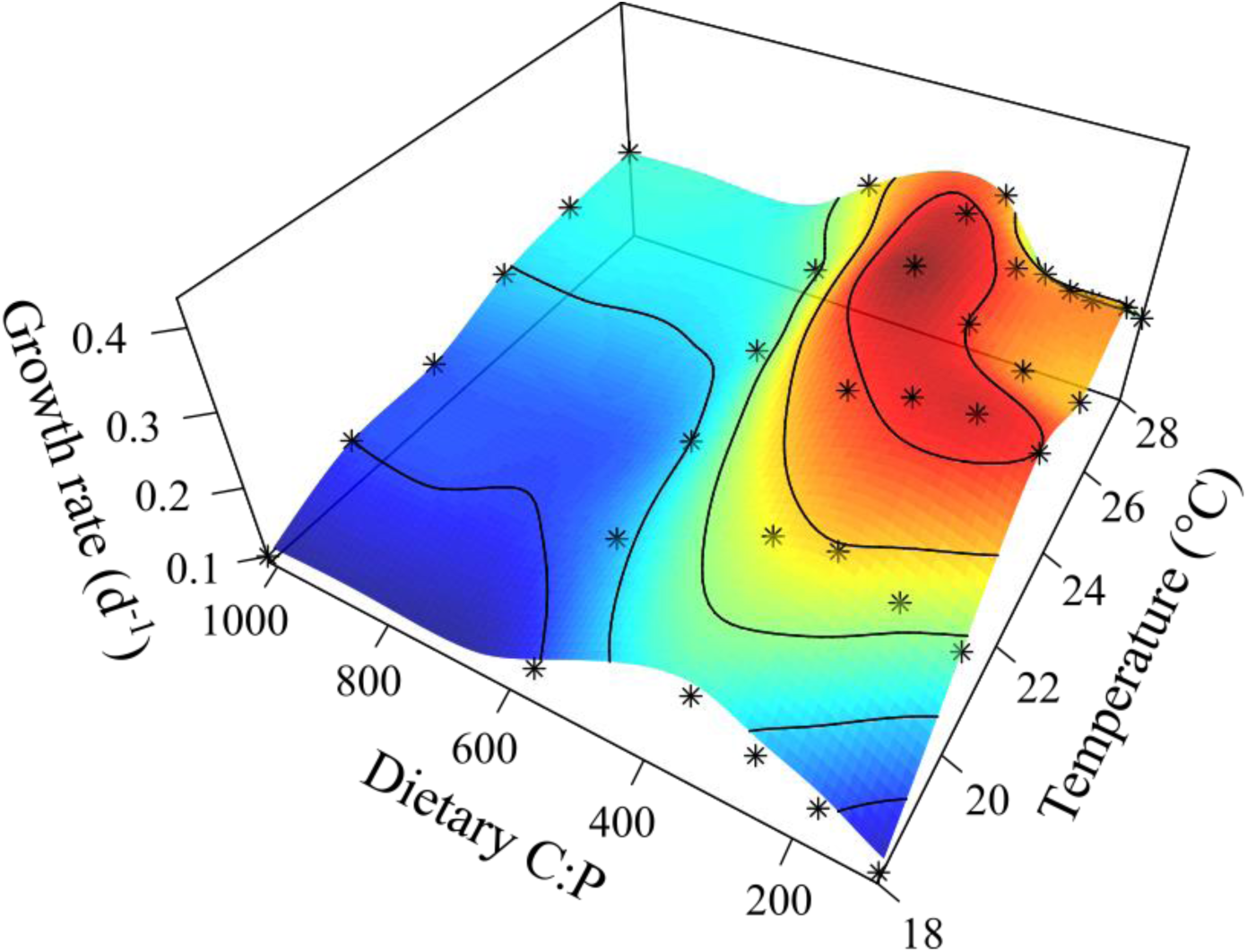
Response surface of *Daphnia manga* somatic growth rate versus dietary C:P and temperature. The asterisks are the experimental observations.

**Figure S3:**
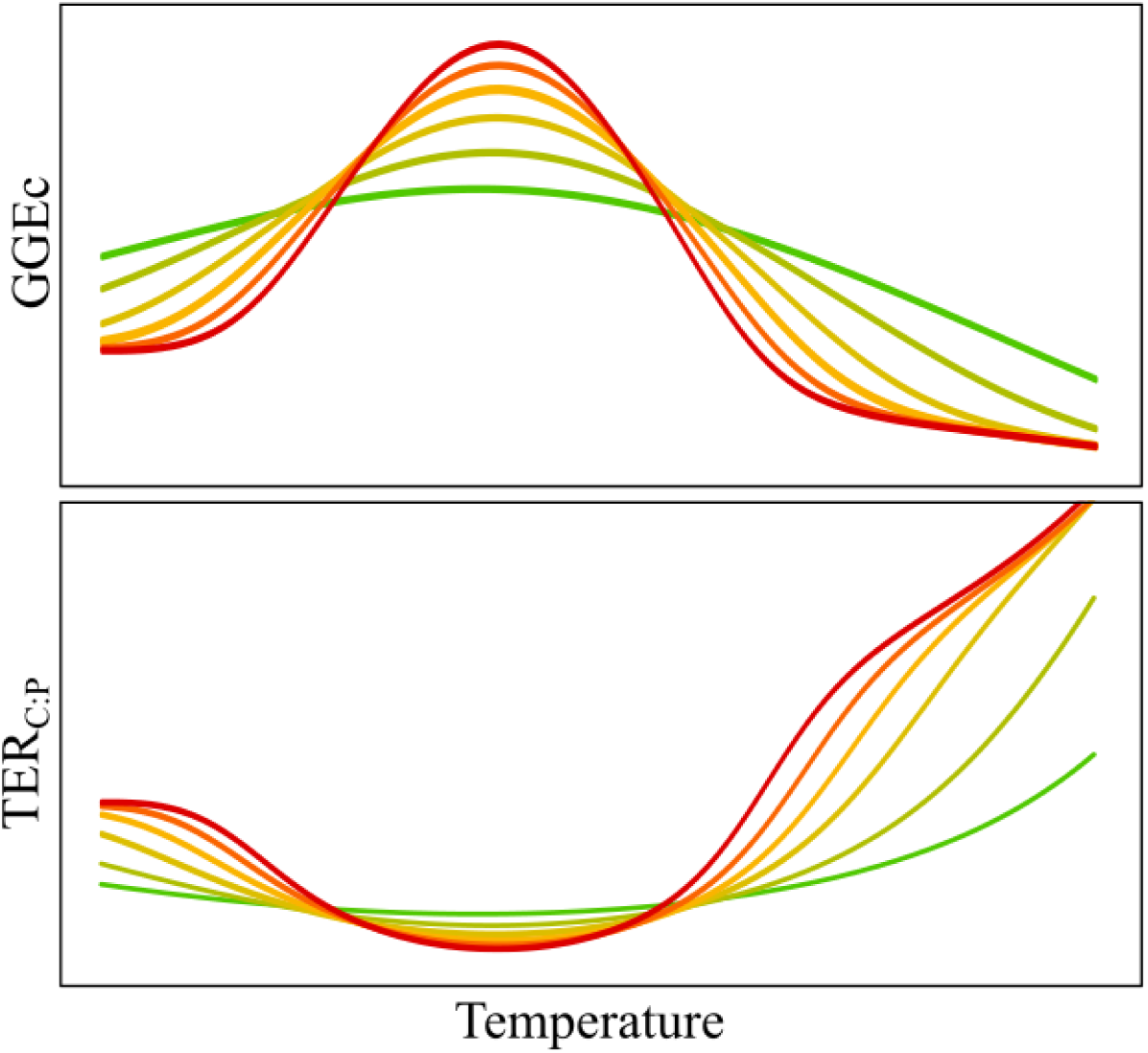
TER_C:P_ reaction norm sensitivity to the variation in the shape of C gross growth efficiency (GGEc) along a specialist (red) –generalist (green) trade-off.

**Figure S4:**
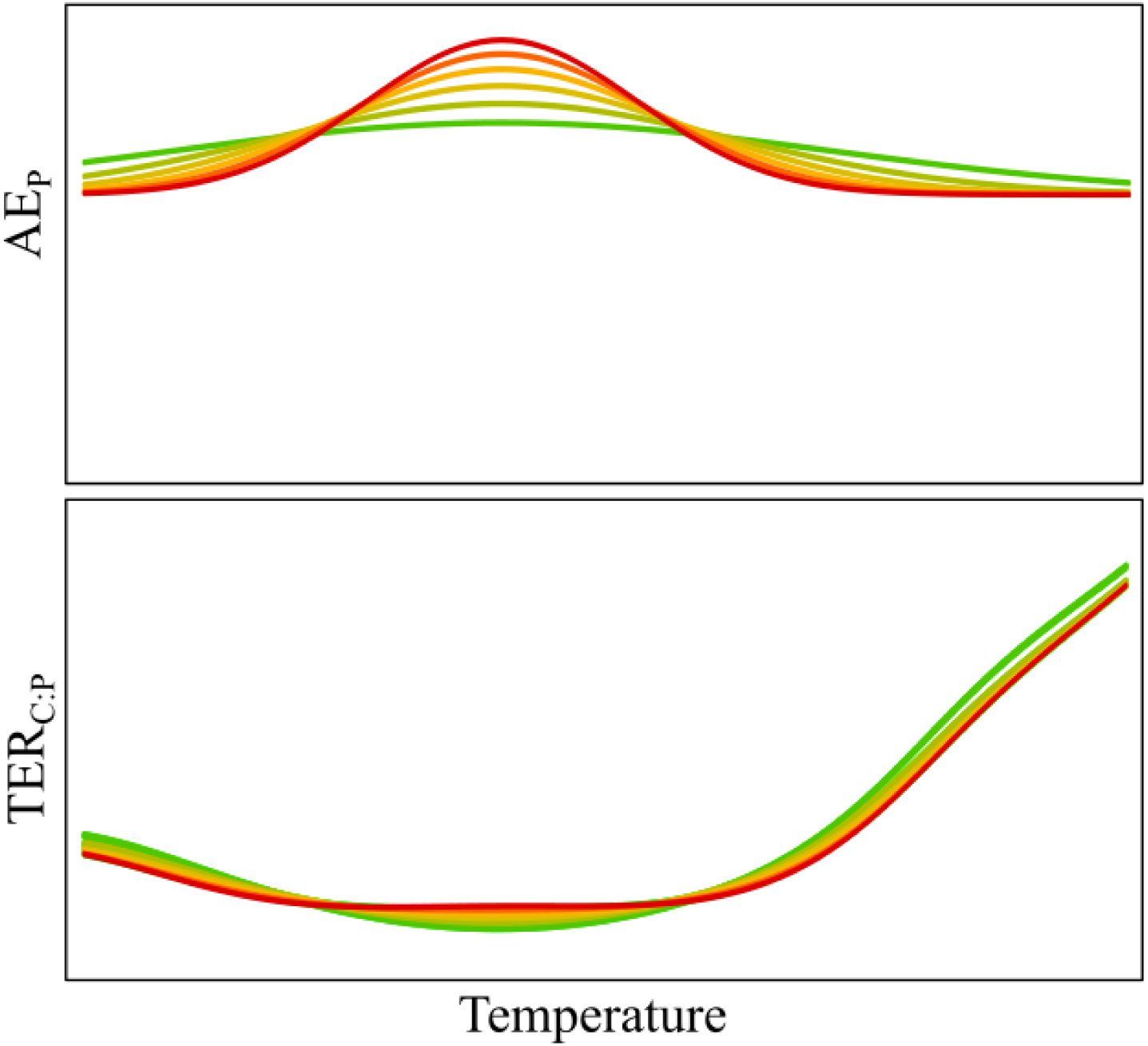
TER_C:P_ reaction norm sensitivity to the variation in shape of P assimilation efficiency (AE_P_) along a specialist (red) –generalist (green) trade-off.

**Figure S5:**
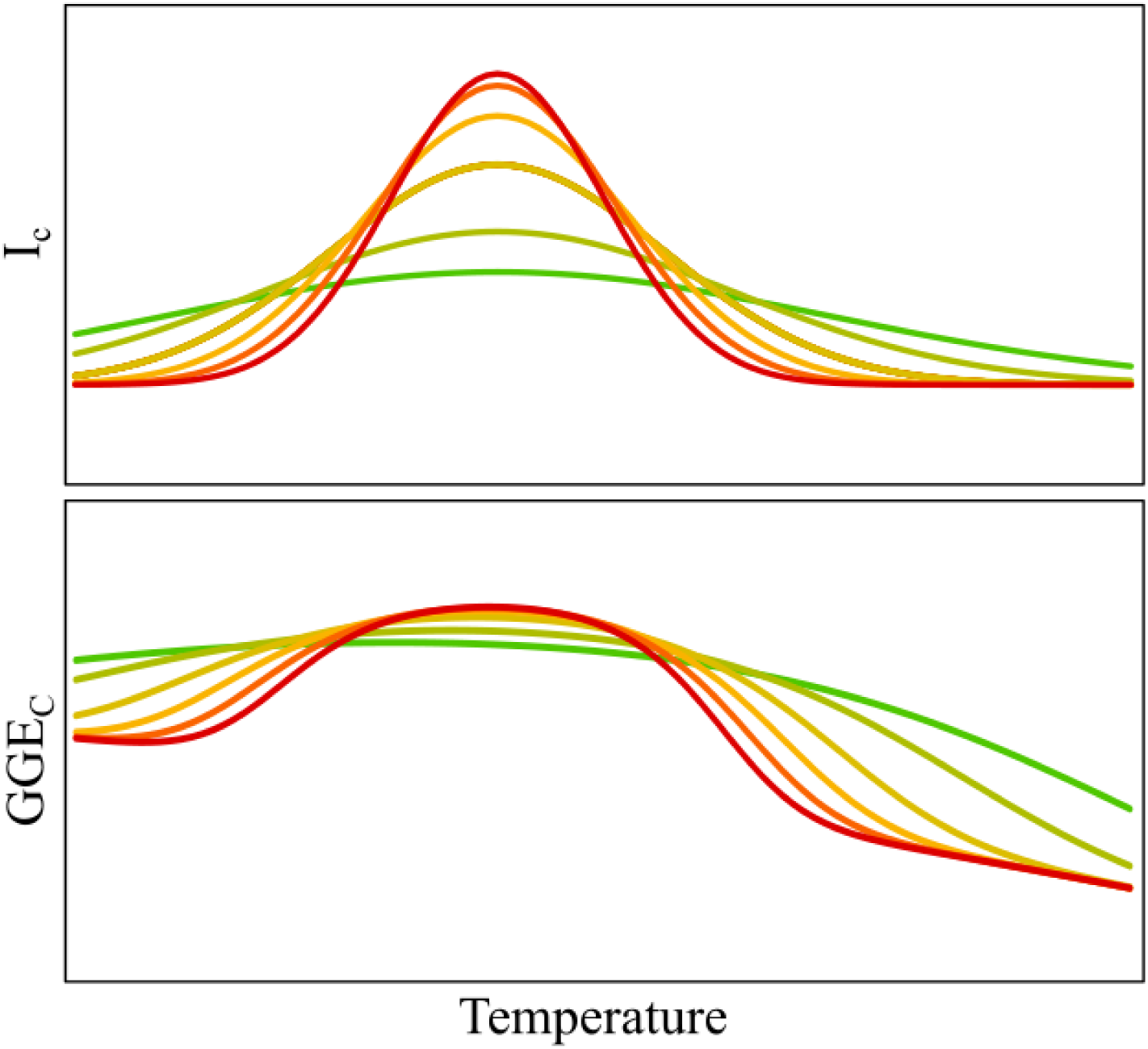
GGEc reaction norm sensitivity to the variation in shape of C ingestion rate (Ic) along a specialist (red) –generalist (green) trade-off.

**Figure S6:**
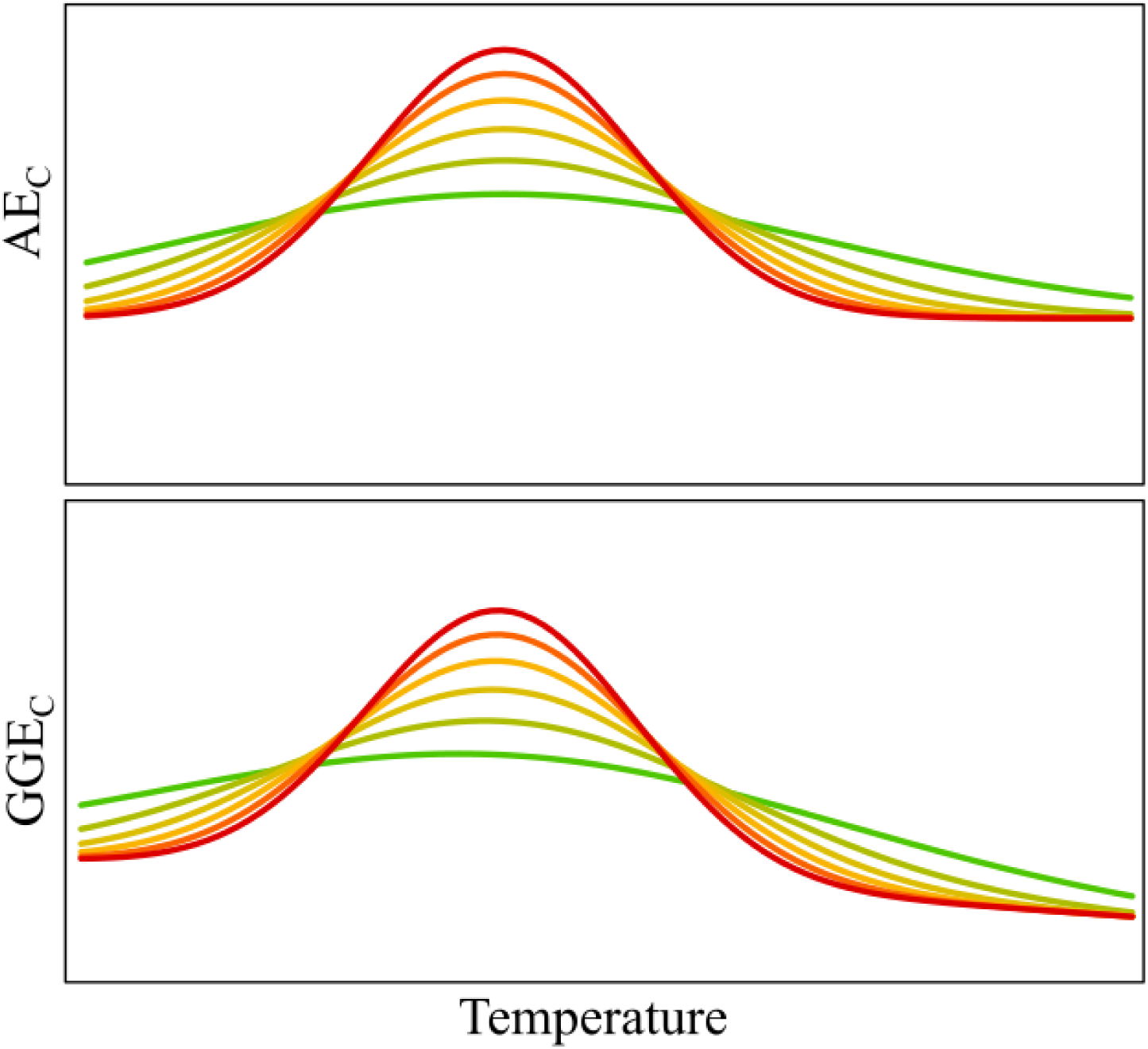
GGEc reaction norm sensitivity to the variation in shape of C assimilation efficiency (AEc) along a specialist (red) –generalist (green) trade-off.

**Figure S7:**
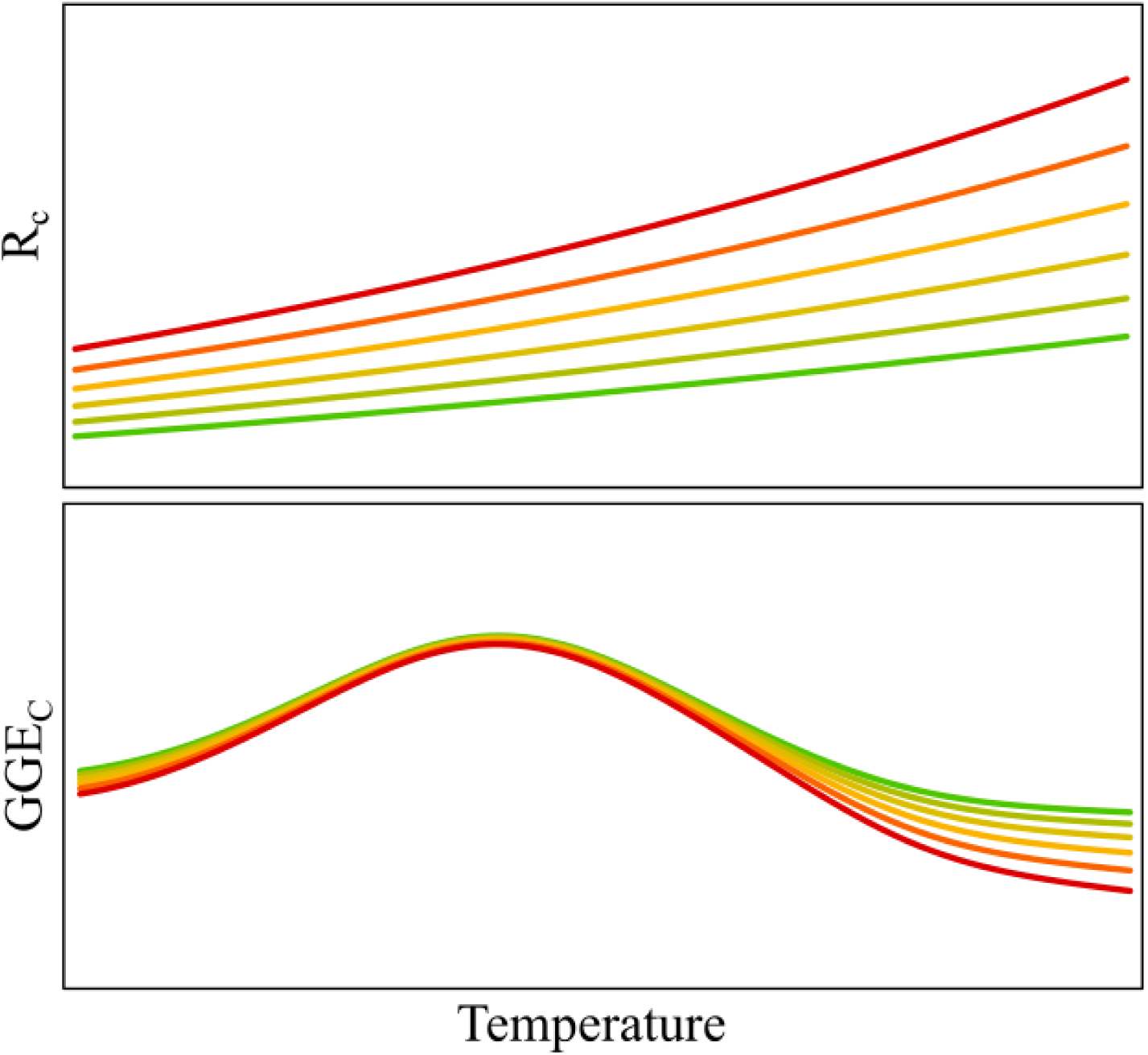
GGEc reaction norm sensitivity to the variation of respiration (Rc) from a slower (green) to faster (red) scaling.

**Table S1:**
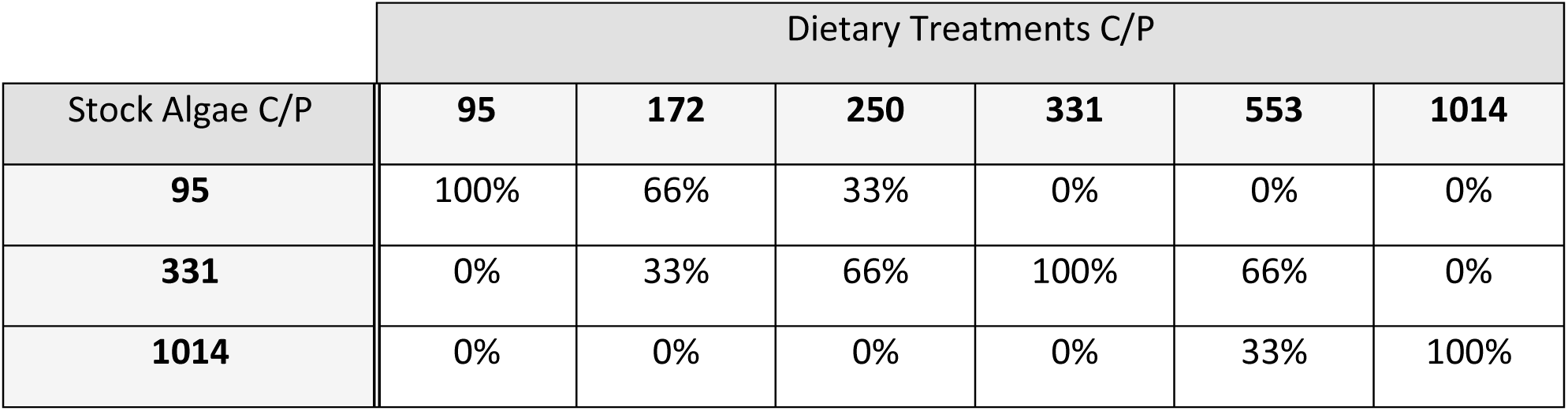
C:P ratio of dietary treatment and proportions of freeze-dried algal stock mixed for their preparation.

|  | Dietary Treatments C/P |  |  |  |  |  |
| --- | --- | --- | --- | --- | --- | --- |
| Stock Algae C/P | 95 | 172 | 250 | 331 | 553 | 1014 |
| 95 | 100% | 66% | 33% | 0% | 0% | 0% |
| 331 | 0% | 33% | 66% | 100% | 66% | 0% |
| 1014 | 0% | 0% | 0% | 0% | 33% | 100% |

